# Does bilingualism buffer genetic predispositions to reading difficulties through alterations of structural interhemispheric connectivity? An ABCD^Ⓡ^ Study

**DOI:** 10.64898/2026.04.07.716864

**Authors:** Marie Lallier, Cristina Rius-Manau, 23andMe Research Team, Amaia Carrión-Castillo

## Abstract

Here, we test the hypothesis that early sustained exposure to complex bilingual environments can positively affect reading development by altering structural interhemispheric connectivity via the corpus callosum (CC). Interhemispheric connectivity has been shown to be inefficient in dyslexia, but also to support compensatory pathways when genetic risk for reading difficulties is present, by enabling the preserved right hemisphere to support a dysfunctional left hemisphere. Mediation models were conducted on children aged 9–10 years (with a 2-year follow-up assessment) from the Adolescent Brain Cognitive Development database (N>10,000). Polygenic scores (PGS) for dyslexia and cognitive performance and continuous bilingualism indices were used as predictors, with reading aloud as the outcome. Bilingualism showed a positive effect on reading partially mediated by the anterior CC, independently of overall brain size. In contrast, genetic predispositions to reading difficulties influenced reading primarily through overall brain size rather than CC connectivity specifically. These two pathways were independent, suggesting that bilingual experience and genetic risk operate through distinct neuroanatomical mechanisms. These findings suggest that recurrent early exposure to complex bilingual environments may shape the brain’s structural connectivity toward a more balanced and integrated bilateral frontal organisation. The results highlight potential brain compensatory pathways induced by environmental experiences that may support more efficient reading development and mitigate risks for developmental dyslexia.

## 1. Introduction

Amongst the cognitive functions that a child must develop during their first years of schooling, reading is by far one of the most complex. Because a child’s brain is highly plastic and adaptable to environmental influences, childhood experiences have a profound impact on shaping and optimising the properties and configurations of neural networks, including those supporting reading development (Dehaene, 2011; Dehaene, Cohen, Morais, & Kolinsky, 2015; Perry, 2002). Reading strongly depends on the adequate development of phonological abilities, which have been a cornerstone theoretical construct in the study of dyslexia (Ramus & Szenkovits, 2008; Snowling M., 1998). Evidence shows that phonological (dis)abilities mediate the link between the complex dyslexia phenotype and its genetic bases (Moll, et al., 2014), and for this reason, they are susceptible to be influenced by linguistic environmental factors and modulate reading outcomes (Landerl, et al., 2013; Snowling M. J., 2008). Therefore, the early acquisition of two phonological systems in early bilingualism might significantly impact the organisation of the reading brain, changing the trajectory of reading acquisition and determining the extent of its success.

Bilingual contexts make early language and phonological acquisition particularly complex, as young learners have to adapt and develop optimal strategies to navigate linguistic variability, uncertainty and conflict. Learning two or more linguistic systems is now widely recognized to lead to neurocognitive adaptations affecting both brain function and structure (Felton, et al., 2017; Pliatsikas, 2019; Pliatsikas, et al., 2020; Amoruso, et al., 2024). Amongst these effects, bilingualism has been associated with reduced classical phonological and attentional control spatial asymmetries or dominance (Hausmann, Durmusoglu, Yazgan, & Güntürkün, 2004; Hull & Vaid, 2006; Hull & Vaid, 2007; Marzecová, Asanowicz, Krivá, & Wodniecka, 2012), indexed by reduced processing advantages for stimuli presented to the contralateral side of the dominant hemisphere, namely the left hemisphere (LH) for phonological operations, and the right hemisphere (RH) for attentional control processes. These neural adaptations might arise because of the need to select and switch between languages, which create perceptual, attentional and cognitive demands that are unique to bilingual language use (Blanco-Elorrieta & Pylkkänen, 2018; Green & Abutalebi, 2013).

Notably, research has shown that tasks with high complexity and attentional demands trigger the activity of LH frontal regions (Friederici, Fiebach, Schlesewsky, Bornkessel, & Cramon, 2005; Roskies, Fiez, Balota, Raichle, & Petersen, 2001; Swick, Ashley, Turken, & U, 2008), involved in both phonological and attentional control networks (Diveica, et al., 2023), and play a critical supporting role when the RH-dominant attentional network reaches maximal engagement (Hirose, et al., 2012). Interestingly, young bilinguals show a hyperreactivity and sensitivity of LH frontal regions to increased attentional demands and complexity (Arredondo, Hu, Satterfield, & Kovelman, 2016; Arredondo, Aslin, & Werker, 2021), which may contribute to shaping a bilateral “phonological attentional control network” ready to handle the complex demands of bilingual environments (Green & Abutalebi, 2013; Kroll & Bialystok, 2013).

This aligns well with observations that, as task complexity increases, a gradual transition is observed going from lateralised and segregated networks towards more integrated or bilateral networks characterised by longer distance and stronger interhemispheric connectivity (Kitzbichler, Henson, Smith, Nathan, & Bullmore, 2011). This hemispheric cooperative benefit is mostly observed in the genu (Davis, Kragel, Madden, & Cabeza, 2011; Davis & Cabeza, 2015), the anterior section of the corpus callosum (CC), a major white matter tract connecting the two hemispheres. These neural connectivity adaptations are further supported by behavioural evidence showing that when a task is sufficiently challenging, hemispheric cooperation is recruited to enhance performance (Belger & Banich, 1998; Hughes, Upshaw, Macaulay, & Rutherford, 2016; Weissman & Banich, 2000), resulting in a “rebalancing” of normally dominant and lateralised networks.

Recently, Klimovich-Gray et al. (2026) proposed a theoretical framework explaining how such rebalancing and strengthening interhemispheric cooperation could underlie successful and resilient neural speech processing adaptations in dyslexia. They suggest that the classically observed bilateral, less left-lateralised, phonological and speech networks in dyslexia - reflecting RH overactivation (Hoeft, et al., 2007; Pugh, et al., 2000)- often viewed as an impairment, may, in some cases, reflect successful compensatory strategies that help manage the high demands imposed by a dysfunctional LH. The authors argue that these successful cases of atypical bilateral neural organisation will partly depend on the active engagement of the CC: an overactivated RH would only serve as an optimal compensatory strategy if its activity is transferred to the challenged LH for support (e.g., (Molinaro, Lizarazu, Lallier, Bourguignon, & Carreiras, 2016; Yu, et al., 2020)). This hypothesis implies explaining why altered structural interhemispheric connectivity - mostly in the posterior splenium CC section, but also the genu - predicts low phonological and reading skills (e.g., (Dougherty, et al., 2007; Frye, et al., 2008; Rumsey, et al., 1996; Swanson, et al., 2015; Robichon, Bouchard, Démonet, & Habib, 2000; Plessen, 2002)) and could be viewed as part of causal accounts of reading deficits.

Here, we adopt an “adaptive” view of the CC where it may either passively suffer from weakened signals sent from a dysfunctional hemisphere - the LH in the case of dyslexia - or support dysfunction through the active (but also more costly) transfer from the preserved hemisphere - the RH in dyslexia. The follow-up task for research is therefore to identify which factor(s) may contribute to favouring the active use of interhemispheric connectivity as an effective compensatory strategy that strengthens the reading networks.

In the present study, we examine early bilingualism as an environmental factor that may ultimately alter interhemispheric connectivity to create a rebalanced and resilient reading network. In line with this, plenty of evidence shows that experience with bilingual environments alters functional and structural brain interhemispheric connectivity mainly in anterior (Bice, Yamasaki, & Prat, 2020; Fedeli, Del Maschio, Sulpizio, Rothman, & Abutalebi, 2021; Luk, Bialystok, Craik, & Grady, 2011; Mohades, et al., 2012; Pliatsikas, Moschopoulou, & Saddy, 2015; Schlegel, Rudelson, & Tse, 2012) but also the posterior (Bice, Yamasaki, & Prat, 2020; Pliatsikas, Moschopoulou, & Saddy, 2015; Pereira Soares, Kubota, Rossi, & Rothman, 2021) CC regions. Moreover, bilinguals seem to exhibit a more balanced and efficient allocation of attentional resources (Bialystok & Craik, 2022; Phelps & Bozic, 2024), which could also reflect the “greater openness” of young bilinguals when they explore their environment (Singh, Kalashnikova, & Quinn, 2023). The CC might have a role to play in this bilingual attentional openness, since it contributes to the optimal attentional orientation across hemifields, thus hemispheres (Chechlacz, Humphreys, Sotiropoulos, Kennard, & Cazzoli, 2015; Pollmann, Maertens, Cramon, Lepsien, & Hugdahl, 2002; Pollmann, 2010), and to reduced lateralisation (Andrulyte, et al., 2024).

This is particularly well illustrated by studies using the dichotic listening paradigm (Kimura, 1961), in which hemispheric dominance for phonological processing and the efficiency of interhemispheric connectivity can be indirectly measured with behavioural readouts. In this paradigm, participants hear different syllables presented simultaneously to both ears and are either instructed to report either the one they heard best (measuring bottom-up attentional orientation) or the one presented in a specific ear (top-down attentional orientation). In this task, right-ear syllables are generally easier to report because they are processed directly by the dominant LH. Left-ear syllables, however, are more complex to report as they must cross through the CC (Pollmann, Maertens, Cramon, Lepsien, & Hugdahl, 2002) from the non-dominant RH to the dominant LH to be processed linguistically (Steinmann, et al., 2017; Steinmann, et al., 2018). Individuals with high degree of bilingual use and exposure have been shown to exhibit increased left-ear reports (Ershaid, 2026; Lallier, Peréz-Navarro, & Ordin, 2024) interpreted as efficient interhemispheric RH-to-LH connectivity. Most importantly this attentional rebalance was associated with better reading and phonological skills in bilingual children and adults reports (Ershaid, 2026; Lallier, Peréz-Navarro, & Ordin, 2024) and with protective effects against family risks of dyslexia in monolinguals (Hakvoort, et al., 2016). This evidence also aligns with reports showing less severe phonological deficits in bilingual individuals with dyslexia (Lallier, Thierry, Barr, Carreiras, & Tainturier, 2018; Lallier, Peréz-Navarro, & Ordin, 2024; Balboni, Kepinska, Berthele, & Golestani, 2025).

Importantly, reading ability is continuously distributed in the population, with dyslexia representing the lower tail of this distribution rather than a categorically distinct condition (Shaywitz, Escobar, Shaywitz, Fletcher, & Makuch, 1992; Pennington, 2006). This dimensional view implies that the neural mechanisms proposed to underlie reading difficulties and associated compensatory strategies in clinically diagnosed individuals, including structural interhemispheric pathways, may be detectable along the continuum of genetic liability in the general population, thus, even below the clinical threshold (Pennington, 2006).

Genetic studies also support a liability model: reading and spelling outcomes and associated cognitive traits (e.g. phonological awareness) in the general population are strongly genetically correlated with dyslexia (Doust, et al., 2022). Genome-Wide Association Studies (GWAS) have begun to identify genetic loci associated with dyslexia (Doust, et al., 2022; Gialluisi, et al., 2020) and reading abilities (Eising, et al., 2022; Price, et al., 2022), enabling the derivation of individual-level polygenic scores (PGS) that aggregate genetic effects into a single genetic predictor for a given trait (Belsky & Harden, 2019; Maier, Visscher, Robinson, & Wray, 2017). PGSs based on the presence of a self-reported dyslexia diagnosis have been shown to predict reading performance (Doust, et al., 2022; Bicona, et al., 2025; Carrion-Castillo, Carreiras, & Lallier, 2025).

Interestingly, PGSs derived from broader and less reading-specific traits, such as cognitive performance (i.e., intelligence) or educational attainment, have been shown to explain a larger proportion of (i) variance in reading outcomes (Procopio, et al., 2024; Carrion-Castillo, Carreiras, & Lallier, 2025) and (ii) associated neural structural organisation (Carrión-Castillo, Paz-Alonso, & Carreiras, 2023), than dyslexia and reading-based PGSs. Cognitive performance PGS effects on reading were found to be mainly mediated by the global brain measure of total left cortical surface area (Carrión-Castillo, Paz-Alonso, & Carreiras, 2023). In addition, a meta-analysis of neuroimaging studies identified smaller overall brain volume as the most robustly replicated structural finding in dyslexia (Ramus, Altarelli, Jednoróg, Zhao, & Scotto di Covella, 2018), an effect that persisted after controlling for IQ. Therefore, it is still unclear whether part of the genetic influence linked to general cognitive performance on reading operates through global brain measures, reading-specific circuitry, or both. Whether PGSs specific to dyslexia also modulate this global brain pathway remains to be established.

Recently, both cognitive performance and dyslexia PGSs were used to investigate how environmental experiences (bilingualism, socio-economic status, etc) modulate genetic influences on reading outcomes. Carrión-Castillo et al. (2025) found a positive effect of bilingualism for reading acquisition that seemed to operate across the genetic risk continuum at the population level, benefiting individuals both at high and low genetic risks. Which brain pathway(s) mediate this positive relationship is still unclear and is the focus of the present study.

Here, we adopt a gene-environment perspective to investigate how genetic predispositions for reading difficulties - linked to less efficiently connected neural systems (Paulesu, et al., 1996; Turker, Kuhnke, Jiang, & Hartwigsen, 2023) possibly including alterations of the splenium and the genu (e.g., (Dougherty, et al., 2007; Sun, et al., 2017) - and bilingualism may both shape the structure of the CC to influence reading outcomes, independently from global brain measures. To do so, we explored the Adolescent Brain Cognitive Development (ABCD^Ⓡ^) database which provides genetic and environmental data of thousands of children across the United States (Jernigan & Brown, 2018). We hypothesised that long-term recurrent exposure to complex dual-language environments would play a key role in molding structural interhemispheric architecture in the brain that is beneficial for learning to read. We expected reading-related PGS and bilingualism to directly influence reading performance, as shown previously, and these associations to be mediated by CC structural variations, with positive (bilingualism) and detrimental (genetic risk, PGS) consequences for reading skills. The structural properties of the genu were expected to be more strongly modulated by bilingualism than reading-related PGS, whereas the splenium was predicted to be more strongly influenced by PGS than bilingualism. Based on previous research in this sample (Carrion-Castillo, Carreiras, & Lallier, 2025), we did not expect the direct effects of bilingualism on reading to strongly interact with genetic risks, but we did not have clear predictions regarding how or whether these two factors would interact to modulate the putative mediating effect of the CC on reading skills (see Figure 1).

**Figure 1.**
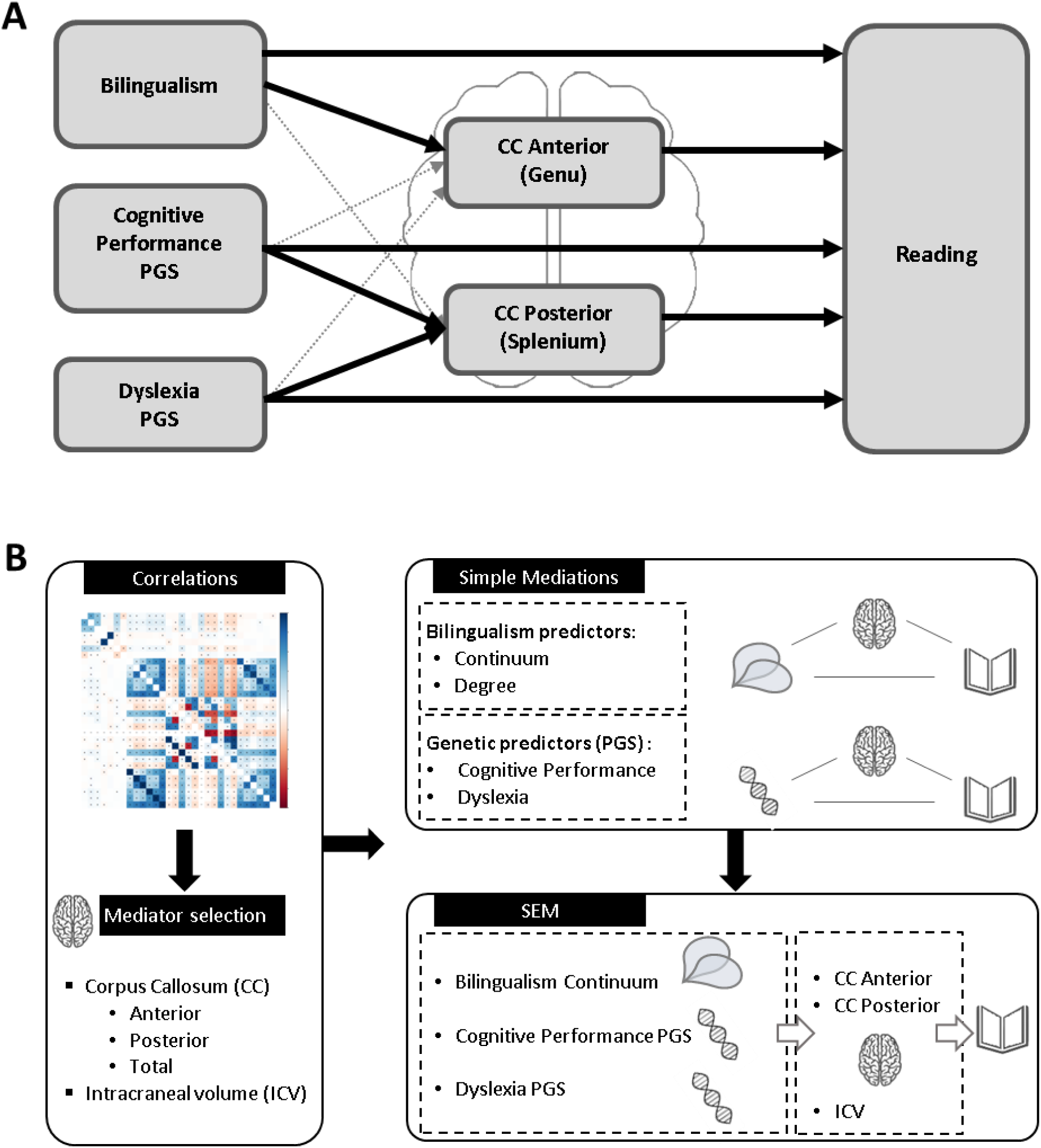
**(A) Conceptual model:** illustrating the hypothesised relationships between bilingualism, polygenic scores (PGS), corpus callosum (CC) structure, and reading outcomes. (**B) Study design workflow.** Summary of the analytical steps and statistical approaches used; see Figure S1 and Table S1 for detailed specifications.

## 2. Methods

### 2.1 Participants

The participants were part of the ABCDⓇ study (https://abcdstudy.org/) (Jernigan & Brown, 2018). At baseline (i.e., first timepoint) they included 11,886 children ages 9 to 10 from the United States recruited and tested between September 2016 and August 2018 (Garavan, et al., 2018). The data was acquired in 21 research centres with the consent of the participant’s parents and following the guidelines of the Declaration of Helsinki. Participants included in the database had data acquired for all the independent variables defined in the experimental design section. Exclusion criteria, including lack of English proficiency, intellectual, medical, neurological or sensory impairments, and absence of first MRI scanning session (Acosta-Rodriguez, et al., 2024).

The current study analysed separately the full baseline sample (N ∼ 11,878) and the 2-year follow-up timepoint from the ABCD Curated Annual Release 4.0. (DOI:10.15154/1523041) and the Genotyping Data from the ABCD Curated Annual Release 3.0 (NDA Study 901; DOI: 10.15154/1519007).

#### 2.1.1 Analysis subsets

The *full sample* included all participants (baseline: N = 11,007, mean age=9.9 years, range=8.9-11; 2-year follow-up: N = 9,693, mean age=12, range=10.6-13.8). Three additional partially overlapping subsets were defined for sensitivity analyses (Table S1): the *full unrelated sample* was derived by retaining one child per family unit (based on rel_family_id), to avoid non-independence due to related individuals (baseline: N = 9,036; 2-year follow-up: N = 7,924). The *European ancestry sample* was defined by restricting to participants within 6 SDs of the European ancestry centroid on the first two genetic principal components (PCs), to minimise population stratification in PGS analyses (baseline: N = 5,740; 2-year follow-up: N = 5,235) (as in (Carrion-Castillo, Carreiras, & Lallier, 2025)). The *European ancestry unrelated sample* combined both filters, retaining unrelated individuals of European ancestry (baseline: N = 4,716; 2-year follow-up: N = 4,286).

Primary mediation analyses for bilingualism predictors were conducted in the full sample to maximise statistical power and generalisability. Primary mediation analyses for PGS were conducted in the European ancestry unrelated sample to minimise population stratification and family-level non-independence.

### 2.2 Variables

All variables included in the current study are listed and described in Table S2, with derived variables defined in Table S3. Extreme outliers were removed for all continuous variables (±7 SD from the mean). Table S4 provides their descriptive statistics per timepoint and subset.

#### 2.2.1 Reading outcome measure

We used the uncorrected reading variable from the NIH Toolbox Oral Reading Recognition Test®, an adaptive test that assesses reading aloud (Gershon, et al., 2014; Luciana, et al., 2018).

#### 2.2.2 Bilingual indexes predictors

Participant bilingualism was quantified using questionnaires completed by both parents and children, covering demographic, acculturation, school, and home environmental factors. While English proficiency was a prerequisite for study participation, questionnaires were administered in both English and Spanish. We operationalized bilingualism through two metrics: a continuous "bilingualism continuum" index and a "bilingual degree" variable for the bilingual subset.

### Bilingualism Continuum Index

This metric reflects the spectrum of linguistic exposure and serves as our main operational definition of bilingualism. We first computed specific sub-scores for proficiency balance, language preference, and environmental exposure from selected questionnaire items (Tables S2, S3). The items incorporated the following dimensions: language proficiency, language usage, linguistic exposure (home and school), parental experience. Then, the score was computed as an absolute weighted average of responses to selected questionnaire items, ranging from 0 (completely monolingual environment) to 4 (maximally bilingual environment). See Table S3 for the specific weighting and scoring for each variable.

### Bilingual Degree Index

This index was derived by recalculating the aforementioned scores exclusively in participants who said they were able to understand or speak a language other than English, thereby excluding monolingual participants.

#### 2.2.3 PGS predictors

The PGS is an individual-level variable, computed from a GWAS study, that represents the weighted summation of variants or single nucleotide polymorphisms (SNPs) associated with the trait multiplied by their regression coefficient as a measure of effect of the SNP within a specific trait (Maier et al. 2018). We used the dyslexia (Doust, et al., 2022) and cognitive performance (Lee, et al., 2018) GWAS summary statistics to compute individual PGS for the target ABCD^Ⓡ^ dataset using PRS-CS (Ge, Chen, Ni, Feng, & Smoller, 2019) (see (Carrion-Castillo, Carreiras, & Lallier, 2025) for details on the procedure).

#### 2.2.4 Brain mediator measures

We used both macrostructural (e.g., volumes) and microstructural (e.g., diffusion tensor imaging) data derived from the ABCDⓇ database’s tabulated data, extracted from a T1-weighted sequence (1 mm isotropic voxels), and a diffusion-weighted MRI sequence, obtained from the scanning sessions in a 3T MRI scanner (General Electric 750, Philips, Siemens) (Casey, et al., 2018).

Macrostructural MRI data were extracted using the standard morphometric pipeline in FreeSurfer 5.3.0, which includes quality control. CC volumes were obtained from the “aseg” segmentation atlas, including anterior, mid-anterior, central, mid-posterior, and posterior segments. Intracranial volume (ICV) was also extracted as part of the “aseg” subcortical atlas to obtain a proxy for overall brain size (Hyatt, et al., 2020).

Diffusion tensor imaging microstructural values were calculated from the CC ROI defined with AtlasTrack (Hagler, et al., 2008) using linear estimation on log-transformed diffusion-weighted signals (Hagler, et al., 2019). Averaged weighted measures included fractional anisotropy (FA), longitudinal diffusivity (LD), transverse diffusivity (TD), and mean diffusivity (MD). Additionally, we included a tractography-derived macrostructural measure of the total CC, namely total fiber bundle volume computed from diffusion-weighted MRI.

### 2.3 Data Analysis

We considered the baseline data for our primary analyses; 2-year follow-up analyses were performed to assess longitudinal consistency and sensitivity of findings. Only participants with complete data on all variables of interest were included in each analysis; no imputation was performed.

#### 2.3.1 Correlation matrices

An exploratory correlation analysis was conducted to inform mediator selection, using Bonferroni correction for multiple comparisons. The selection criteria were: (i) significant correlations with reading outcomes, and (ii) for variables meeting this criterion, theoretical relevance to prioritise among candidates from the same structure (i.e. CC subregions).

#### 2.3.2 Mediation analyses

We first performed mixed effects models using the *lme4* package (Bates, Mächler, Bolker, & Walker, 2015) to define direct effects between each predictor and outcomes.

Then, we conducted simple mediation analyses using the *mediation* package (version v4.5.0) (Tingley, Yamamoto, Hirose, Keele, & Imai, 2014), with 10,000 quasi-Bayesian simulations. In all models, age and sex were included as fixed-effect covariates and site as a random effect to account for the nested data structure. To control for global brain size, ICV was included as a covariate in both the mediator and outcome models. For the PGS analyses, genetic ancestry PCs were included as additional covariates to account for population stratification (Patterson, Price, & Reich, 2006). These PCs were derived based on genotype data within each full and European ancestry subsets separately (see (Carrión-Castillo, Paz-Alonso, & Carreiras, 2023)). To control for family-wise error rate across the 48 mediation tests (4 predictors × 4 mediators × 3 outcomes), Bonferroni correction was applied. The threshold for statistical significance was adjusted to α = 0.05/48 = 0.001. Results are presented with both raw and corrected p-values in Tables S5-S8.

All sensitivity analyses are specified in section 2.3.4 and Table S1. In short, to assess robustness to global brain size adjustment, secondary mediation analyses were repeated with CC mediator measures normalised by ICV, and without ICV adjustment (Dick, et al., 2021). Family-level relatedness could not be modelled as an additional random effect in the mediation; to address this, analyses were repeated in “*unrelated”* subsets, and a sensitivity analysis incorporating family as a random effect (instead of site) was conducted for the primary finding using *lme4* directly. Sensitivity analyses were not corrected for multiple comparisons given their exploratory nature, but are reported for transparency.

Analyses were replicated with *lavaan* (v0.6-15) (Rosseel, 2012) with the MLR estimator with cluster-robust standard errors, and study site as a cluster variable, to provide a bridge to the structural equation modelling framework described below. The *mediation* package results are reported as primary given its more complete handling of the nested data structure through mixed-effects models than the *lavaan* models.

#### 2.3.3 Structural equation modelling (SEM)

To extend the simple mediation analyses and examine all predictors and mediators simultaneously, we fitted a near-saturated SEM in *lavaan* (Rosseel, 2012). This model was fit for the baseline timepoint and included all predictor and mediator variables (except the “bilingual degree” variable, which was not significant in any of the mediation models) and reading as outcome. ICV was modelled as an endogenous variable predicted by all three predictors, and was additionally included as a covariate in the anterior CC and posterior CC equations. Residual covariance between anterior CC and posterior CC was freely estimated to account for shared variance between adjacent subregion measures.

The model was estimated using the MLR estimator with standard errors robust to non-normality. Indirect effects were tested using the delta method approximation, as bootstrap confidence intervals cannot be combined with cluster-robust standard errors in *lavaan*. Age and sex were included as covariates in all equations, and study site was specified as a clustering variable to obtain cluster-robust standard errors. No additional correction for multiple comparisons was applied to SEM path coefficients, as the model was estimated simultaneously and serves to confirm patterns observed in the primary mediation analyses rather than test independent hypotheses. Robustness checks were run with additional covariates (Table S1, Figure S1, section 2.3.4 S*ensitivity analyses*).

Model fit was evaluated using SRMR, which remains interpretable in near-saturated models (df=2). CFI, RMSEA, and TLI are not reported as they are known to perform poorly with very low degrees of freedom (df = 2) (Kenny, Kaniskan, & McCoach, 2014) and produce uninterpretable values in the current model. Model adequacy and robustness of findings were therefore primarily evaluated based on SRMR and the consistency of parameter estimates across model specifications.

#### 2.3.4 Sensitivity analyses (Table S1)

Sensitivity analyses examined robustness across the following dimensions:

1. Sample composition: all analyses were repeated in four subsets differing in ancestry stratification and relatedness filtering (Tables S1, S4). Primary results on bilingualism effects on reading are reported for the full sample, and for PGS effects for the European unrelated sample. The SEM was conducted in the full sample to retain sufficient power when modelling bilingualism and PGS effects simultaneously.
2. Given evidence that covariate selection can substantially alter structural-behavioral associations (Hyatt, et al., 2020), we examined the robustness of our findings across three analytical approaches: unadjusted models (NOadj), models with ICV as a covariate (ICVcov), and models using brain volume normalized by ICV (ICVnorm).
3. Analytic approach: mediation analyses were replicated using *lavaan* to confirm robustness to estimation framework (see Section 2.3.2).
4. Ancestry PCs and SES: SEM models were re-run adding ancestry principal components (PC1–PC10) and socioeconomic covariates (household income, parental education).
5. To assess specificity of findings to reading, analyses were repeated with control cognitive outcomes vocabulary (NIH Toolbox Picture Vocabulary Test) (Gershon, et al., 2014) and non-verbal reasoning (WISC-V Matrix Reasoning) (Wechsler, 2014) as outcomes (Luciana, et al., 2018).

A full overview of all analysis specifications is provided in Table S1.

The main hypotheses of this study were preregistered in OSF: https://osf.io/hcf53 (Rius-Manau, Lallier, & Carrion-Castillo, 2026). Deviations from the pre-registered analyses are detailed in the Supplementary materials (Supplementary annex A).

## 3. Results

Descriptive statistics for all variables and subsets are presented in Table S4. The results of the correlation analyses (Figures S2 and S3) led to the selection of anterior and posterior CC volumes and ICV, and the total CC fiber bundle volume (diffusion-weighted MRI).

### 3.1 Direct effects of the predictors on reading (ADE paths in Tables S5-S8)

*<u>Bilingualism → Reading:</u>* Bilingualism continuum scores were positively and robustly associated with reading across all timepoints (baseline std. β ≈ 0.033-0.037, unadjusted p < .0001; year 2 follow up: std. β ≈ 0.0548–0.0575, unadjusted p < .0001; Table S5) and subsets (Tables S6-S8). The bilingualism degree score computed within the bilingual subsample only (N = 3,767 at baseline and year 2 follow up: N=2,654) showed nominal associations with reading in some subsets and timepoints that did not survive correction for multiple comparisons (Table S5-S8).

Regarding the control outcomes (vocabulary and non-verbal reasoning), bilingualism continuum showed no significant effect on non-verbal reasoning in any subset, and only a nominally significant effect on vocabulary in the full European ancestry subset at baseline that did not replicate in the unrelated subset. A negative association between bilingualism degree and non-verbal reasoning was nominally significant in the full samples with bilingualism degree scores (std. β = −0.035 to −0.045, p = .009–.032, N=3,735 to 3,133), but was absent in European ancestry subsets.

#### PGSs → Reading

Both PGSs showed strong and consistent effects in all subsets (Tables S5-S8): CP PGS was positively (std. β ≈ 0.203-0.263, all unadjusted p < .0001), and Dyslexia PGS was negatively (std. β ≈ −0.145 to −0.195, all unadjusted p < .0001) associated with reading.

Regarding the control outcomes, CP PGS also showed strong and consistent positive effects on both vocabulary (std. β = 0.15-0.223, p < .0001) and non-verbal reasoning (std. β = 0.12-0.148, p < .0001) across subsets and timepoints. Dyslexia PGS effects showed mostly nominally significant negative effects on vocabulary in the European ancestry subsets at baseline and follow-up (std. β = -0.058 - -0.011, p < .0001–0.2548) but weaker effects for non-verbal reasoning which did not survive in the European unrelated subset (Table S8).

### 3.2 Mediation analyses

Full results are presented in Supplementary Tables S6–S8 and in Figure 2. Point estimates showed near-perfect agreement across methods (92.5% agreement on statistical significance of indirect effects; Supplementary Annex B), supporting the robustness of findings across estimation frameworks (Table S9).

**Figure 2.**
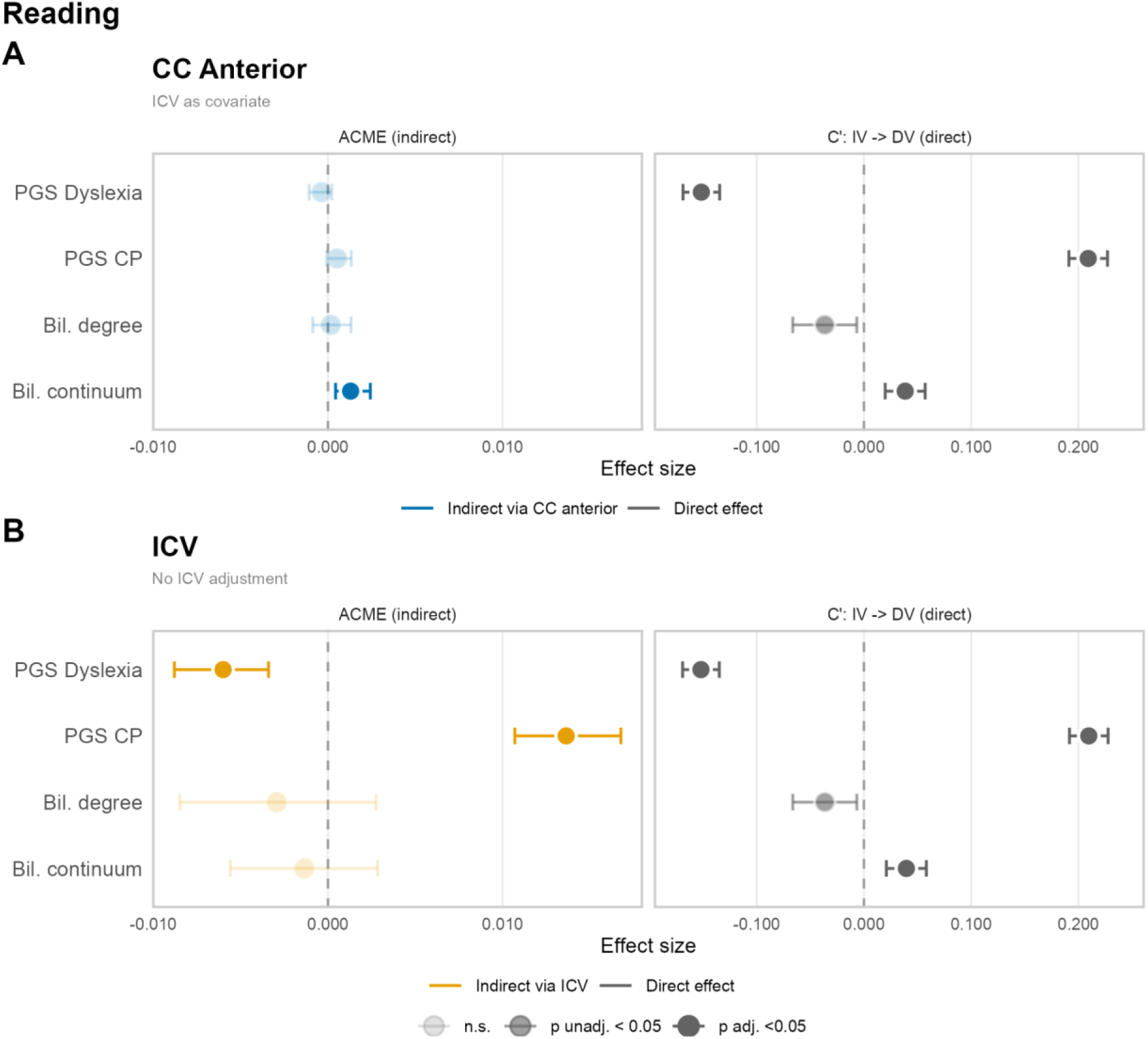
Forest plots showing direct and indirect effects of bilingualism and polygenic score on reading. Mediations are shown for **(A)** anterior corpus callosum (CC) and **(B)** intracranial volume (ICV). Results are shown for the full sample at baseline (maximum N =10,708, varying by analysis due to missing data), using the mediation package. Note that primary PGS analyses were conducted in the European ancestry unrelated subset (maximum N = 4,636); results for this subset are reported in Supplementary Figure S5, Table S8. Effect sizes are standardised (outcome and mediator z-scored prior to model fitting). Faded points indicate non-significant effects (p ≥ .05, unadjusted); error bars represent 95% bootstrap confidence intervals (10,000 simulations, not necessarily symmetric around the point estimate). Sensitivity analyses across ICV specifications and sample subsets are reported in Supplementary Figure S5.

#### 3.2.1 Mediation analyses for the prediction of bilingualism to reading

There was no significant mediation for bilingualism degree (only bilingual participants) (Table S5, Figures S4, S5), which showed high kurtosis across subsets (Table S4). For bilingualism continuum (including both monolinguals and bilinguals), analyses revealed a significant indirect effect on reading via the anterior CC volume at baseline, before and after ICV adjustment (Figure 2A, Table S5). Specifically, after including ICV as a covariate, bilingualism continuum was negatively associated with anterior CC volume (std. A-path = - 0.0319, p = 0.007), and anterior CC volume was negatively associated with reading (std. B-path = -0.0354, p<0.0001), yielding a significant positive indirect effect (std. indirect β = 0.0013, 95% CI [0.0004, 0.0024], p = 6x10-4) (Table S5). Strikingly, this indirect effect was *negative* and significant without ICV control (std. indirect β =-0.0018, 95% CI [-0.0031, - 0.0007], p<0.0001; std B-path = 0.0511, p<0.0001), reflecting the zero-order negative correlation between bilingualism and anterior CC volume (r = -0.04, p=3x10-5, Figure S2) and leading to a reversed effect after ICV was partialled out. This indirect effect remained significant across ICV handling specifications, and when family ID was modelled as a random effect instead of study site (p = 0.048 to <0.0001, Table S10).

Robustness analyses in other subsets across the two timepoints showed that this indirect effect was not significant at the 2-year follow-up (mean age 12, Table S4) in neither subset (Tables S5-S8; with ICV adjustment: full sample std. indirect β =0.0004, p =0.176, N=6,796). However, it was maintained in the unrelated full sample at baseline (N = 8,840, Table S6), with both ICV-covariate and ICV-normalised specifications, but not after adjusting for ancestry PCs (Table S9, Figures S5-S6). In the full European ancestry subset at baseline (N = 5,389, Table S7), this indirect effect was nominally significant with ICV as a covariate (std. indirect β = 0.0011, p = .034) and with ICV normalisation (std. indirect β = 0.00117, p = .014). In the unrelated European subset (N = 4,635, Table S8), no significant indirect effect was observed (all p > .10). The full robustness analysis of the indirect effect of bilingualism on reading through the anterior CC across 36 model specifications is presented in Figure S6.

### Control outcomes (vocabulary and non-verbal reasoning)

There were no significant mediations for bilingualism degree. Anterior CC (unadjusted for ICV or ICV-normalised) significantly mediated the effect of bilingualism on non-verbal reasoning and vocabulary (Table S5), but not when including ICV as a covariate (Figures S7A, S8A). These effects were not present in the European ancestry subsets (Tables S7,S8).

#### 3.2.2 Mediation analyses for the prediction of PGS to reading

##### PGS→CC →Reading

The total CC fiber bundle volume showed significant indirect effects for both PGS (CP PGS: std. indirect β = 0.0068, p <0.0001; Dyslexia PGS: std. indirect β = - 0.005, p <0.0001, Table S8). However, these indirect effects were no longer significant after including ICV as a covariate or after ICV-normalised CC volume (all p > .05). This pattern was consistent across subsets (Tables S5-S8, S9). There was a nominally significant indirect effect of the PGS on reading through the anterior CC (baseline ICV-normalised: p=0.026) that was affected by ICV handling and subset specifications (Tables S5-S9).

##### PGS→ICV→Reading

Both CP PGS and Dyslexia PGS showed significant indirect effects on reading via ICV. Specifically, higher CP PGS was associated with larger ICV (std. A-path at baseline= 0.0868, p = 4.1x10-13; year 2=0.0809, p=5.1x10-08), which in turn was positively associated with reading (std. B-path at baseline= 0.1516, p = 1.4x10-19; year 2= 0.1646, p=2.8x10-16), yielding a significant positive indirect effect (std. indirect β at baseline= 0.0131, 95% CI [0.0089, 0.0179], p<0.0001; year 2=0.0133, 95% CI [0.0079, 0.0194], p<0.0001). Conversely, higher Dyslexia PGS was associated with smaller ICV (std. A-path at baseline = -0.0435, p = 0.0003; year 2=-0.0355, p=0.0165), yielding a significant negative indirect effect on reading via ICV (std. indirect β at baseline=-0.0074, 95% CI [-0.0117, - 0.0033], p = 0.0004; year2=-0.0069, 95% CI [-0.0124, -0.0012], p=0.0148). These patterns were consistent across all subsets (Tables S5-S9) and timepoints (Figure S4).

#### Control outcomes (vocabulary and non-verbal reasoning)

CP PGS showed significant total effects and ICV-mediated indirect effects for vocabulary (Figure S7C, Table S8) and non-verbal intelligence (Figure S8C, Table S8). Dyslexia PGS total effects were nominally significant, though marginal ICV-mediated indirect effects were observed for both (vocabulary: std. indirect β at baseline =-0.007, p =0.0004, year 2=-0.0062, p<0.0194 ; non-verbal reasoning: std. indirect β =-0.0048, p=0.0002) (Table S8, Figures S7C, S8C).

### 3.3 Structural equation modelling (SEM)

The model is presented in Figure 3 including all predictors and mediators tested in the previous analyses, except for bilingual degree that was not significant in any of the models. Overall, the model fit for the primary SEM was acceptable: SRMR = 0.059. CFI and RMSEA are not interpreted given the near-saturated model structure (df = 2, see Methods). R² values for the endogenous variables were: ICV = 0.24 (variance explained by PGS and bilingualism predictors), CC anterior = 0.03 and CC posterior = 0.03 (variance explained by predictors and ICV), and reading = 0.16 (variance explained by all predictors and mediators).

**Figure 3.**
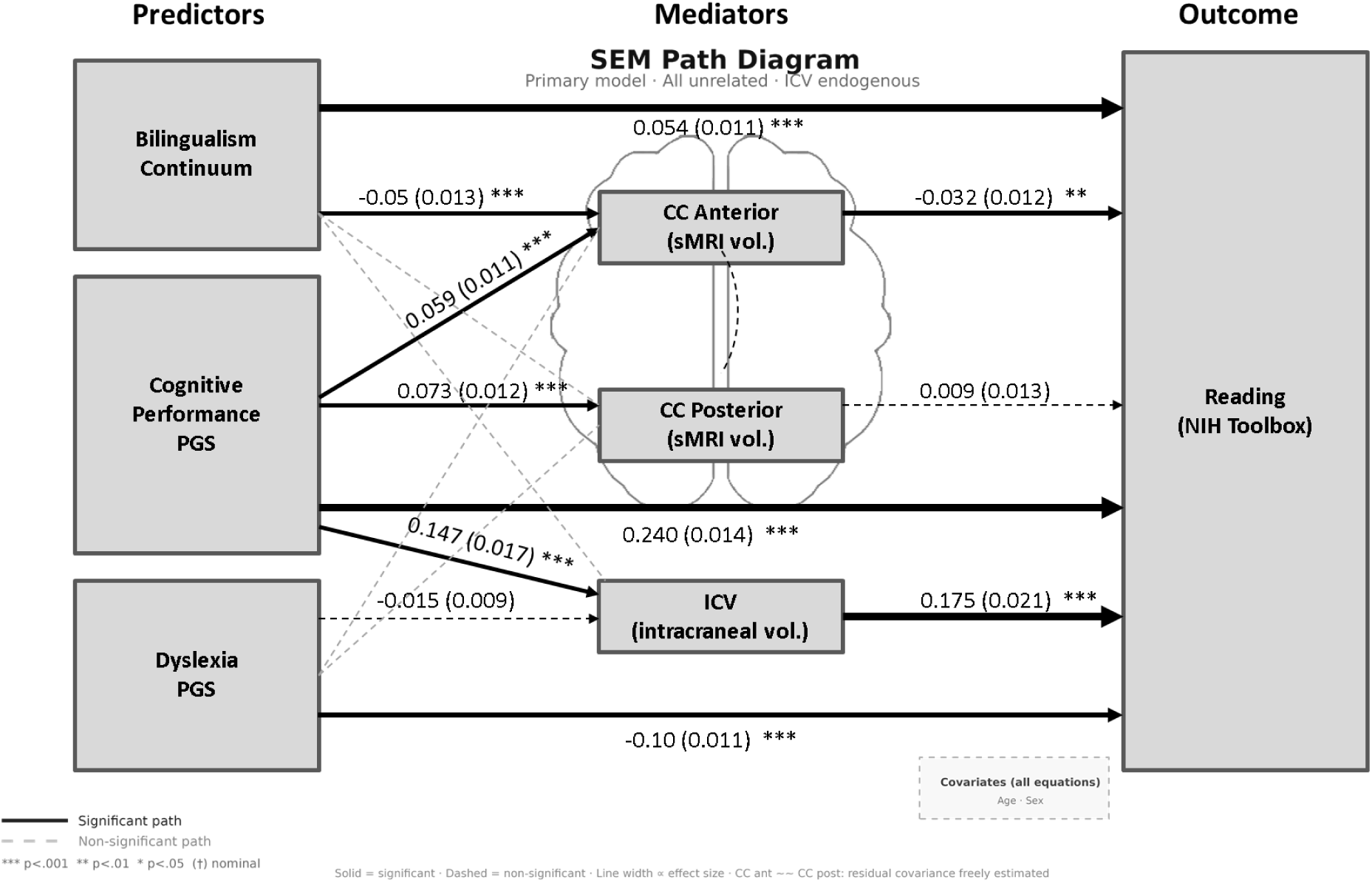
Structural equation model path diagram, full sample. Results shown for the full sample at baseline (N = 9,634); full results in Table S11. Nodes represent observed variables; paths show standardised coefficients (standard error). Solid lines indicate significant paths (p < .05); dashed lines indicate non-significant paths. Line width is proportional to effect size. Model estimated using MLR with cluster-robust standard errors (site as clustering variable). ICV was modelled as an endogenous mediator for polygenic score effects and as a covariate in CC subregion equations (ICV → CC paths not shown for clarity); CC anterior and CC posterior residual covariance was freely estimated (∼∼). All equations included age and sex as covariates.

#### Direct effects

The direct effects of the bilingualism and PGS predictors on reading were stable across SEM model specifications with and without ICV, and additional covariate adjustment (e.g. CP PGS std. β=0.16-0.26, Dyslexia PGS std. β= -0.10 - -0.116; bilingualism continuum std. β =0.040-0.064; full results in Table S11).

### Indirect effects

#### Bilingualism→CC→Reading

The indirect effect of bilingualism via anterior CC reported in the simple mediation analyses was nominally significant in the full sample (std. indirect β (SE) = 0.002 (0.001), p = .0346), also after adjusting for SES covariates (Table S11). However, it disappeared with ancestry PC inclusion (p = .26).

#### PGSs→CC→Reading

The CP PGS showed a nominally significant indirect effect via anterior CC in the model without ancestry PCs (std. β = −0.002, SE = 0.001, p = .017), but this effect did not survive PC adjustment. No significant indirect effect via posterior CC was observed.

#### PGSs→ICV→Reading

In the primary model (with ICV and no ancestry PCs), the CP PGS showed a robust indirect effect on reading via ICV (std. β = 0.026, SE = 0.004, p = <0.001). This effect was stable across all model specifications. The Dyslexia PGS indirect effect via ICV was not significant in the primary model (std. β = −0.003, SE = 0.001, p = .072) but was nominally significant on all other specifications (Table S11).

*Control outcomes (vocabulary and non-verbal reasoning)* (Tables S12 - S13).

Analyses confirmed robust direct effects of CP PGS and significant indirect effects via ICV (vocabulary: std. β = 0.026, SE=0.005, p <0.001; WISC-V: std. β = 0.018, SE=0.0028, p <0.001). The direct effect of Dyslexia PGS, and indirect effect through ICV were nominally significant for some model specification only (Tables S12, S13). Additionally, indirect effects of both PGS via posterior CC emerged in these models, strongest for vocabulary, but also present for non-verbal reasoning (vocabulary: std. β = 0.003, SE=0.0009, p <0.001; WISC-V: std. β = 0.003, SE=0.001, p = 0.012). This indirect effect was attenuated but present after adjusting for ancestry PCs, ICV and SES (Tables S12, S13).

## Discussion

By integrating genetic, brain and behavioural data, we attempted to provide a possible mechanistic understanding of how the early and recurrent exposure to bilingual environments may sharpen the resilience of the brain structure to genetic predispositions to reading difficulties. Our hypothesis more specifically focused on the CC, which has been proposed to hold a critical role in contributing to the formation of a less lateralised and rebalanced resilient reading network (Klimovich-Gray, Bozic, Molinaro, & Lallier, 2026).

### 1. Global brain size (but not the CC) mediates the influence of genetic predispositions to reading difficulties

Both PGS predicted reading performance (negatively for Dyslexia PGS and positively for CP PGS), as reported in previous studies (Carrión-Castillo, Paz-Alonso, & Carreiras, 2023; Carrion-Castillo, Carreiras, & Lallier, 2025). Here, we showed that this effect was present in the full subsets, which included heterogeneous ancestries, as the ABCD study is diverse by design, aiming to obtain a representative sample from the general US population (Garavan, et al., 2018). Effect sizes were somewhat larger in the European ancestry subsets, consistent with the known reduction in PGS predictive validity in ancestrally diverse samples due to GWAS discovery bias (Duncan, et al., 2019; Martin, et al., 2017), though the overlap in intervals suggests robustness to ancestry composition.

These direct effects were found to be mediated by the total CC volume before adjusting for ICV, but not after ICV controls. This reflects shared variance between CC volume mediation and global brain size, rather than a neurogenetic interhemispheric pathway to reading outcomes. Indeed, part of both the CP and the Dyslexia PGS direct effects on reading were mediated by a positive effect of ICV on reading, although less consistent and smaller in magnitude for Dyslexia PGS. Crucially, these indirect effects were robust after controlling for socioeconomic background and to analytical choices, and were found for both study timepoints. These findings are in line with the consistent overall smaller ICV measures in dyslexia (Ramus, Altarelli, Jednoróg, Zhao, & Scotto di Covella, 2018), as well as with the indirect effect of CP PGS on reading through total LH cortical surface area in this sample (Carrión-Castillo, Paz-Alonso, & Carreiras, 2023). Interestingly, the SEM analyses revealed that the direct effects of both PGS and indirect effects through ICV on reading remained partially independent when estimated simultaneously. Overall, these results suggest that both reading-related PGS have an influence on ICV (positive for CP, negative for Dyslexia) which in turn robustly modulates reading-specific and non-reading-specific cognitive outcomes (with some variation between SEM and mediation analyses for control outcomes). Notably, an indirect pathway from Dyslexia PGS through ICV more specific to reading might capture variance in core processes underlying dyslexia, such as grapheme-to-phoneme mappings, independent of oral vocabulary and nonverbal IQ.

Contrary to our predictions, there was no robust indirect effect of PGS on reading through the CC, as effects disappeared when controlling for ICV. This suggests that, rather than deficit-based neurogenetic mechanisms, the previously reported links between CC and dyslexia (e.g. (Vandermosten, Poelmans, Sunaert, Ghesquière, & Wouters, 2013)) may reflect compensatory strategies that we hypothesise are promoted by complex experiences, such as exposure to bilingual environments.

### 2. The positive effects of bilingualism on reading are partly mediated through the anterior CC, independently of global brain size and genetic predispositions to reading difficulties

We found a highly robust direct positive effect of our bilingualism-continuum index on reading - including both monolingual and bilingual children (note that this effect was absent in smaller samples composed solely of bilingual participants) - across both time points, in both mediation and SEM analyses, replicating the findings of Carrión-Castillo et al. (2025) while using a continuous rather than dichotomous operationalisation of bilingualism: the greatest reading benefits were seen in children exposed to the most bilingual environments. This replicates previous behavioral findings in Basque–Spanish Grade 1 bilinguals (Lallier, Peréz-Navarro, & Ordin, 2024), showing that bilinguals predominantly exposed to dual-language contexts demonstrated more advanced (lexical) reading skills than bilinguals mainly exposed to single-language environments. This highlights the importance of taking into account the heterogeneity of bilingual experiences to characterise their effects on neurocognitive outcomes (see (Blanco-Elorrieta & Pylkkänen, 2018; DeLuca, Rothman, Bialystok, & Pliatsikas, 2020).

Although a priori planned, we chose not to directly test the interaction between PGS and bilingualism since they predicted reading through independent neural paths (ICV for PGS, CC for bilingualism). Of note, Carrión-Castillo et al., (2025) found a weak nominal genes x environment interaction in this ABCD sample which tended to show stronger protective effects for bilinguals with the highest risks of developing reading difficulties. In any case, these findings suggest that learning multiple languages boosts reading development across the whole risk continuum.

Critically and as predicted, the direct effect of bilingualism on reading was mediated by the CC, its anterior region specifically, in the main analysis using the full sample. This finding supports our hypothesis that early exposure to complex bilingual environments promotes the active engagement of frontal interhemispheric connectivity to create a rebalanced attentional control network. Importantly, this effect was robust despite controlling for global brain size (also in both European samples), and was relatively specific to reading, especially in the mediation analyses. This indirect effect was attenuated with ancestry adjustment, smaller sample sizes, and more conservative estimation approaches (e.g., only nominally significant and less specific to reading in the SEM analyses), calling for replication in other bilingual populations in which linguistic diversity and genetic ancestry are less confounded (see (Ershaid, 2026; Lallier, Thierry, Barr, Carreiras, & Tainturier, 2018; Lallier, Peréz-Navarro, & Ordin, 2024)).

The selective mediation of the anterior (not posterior) CC independently of ICV rules out a general whole-brain structural origin. This specific mediation aligns with frontal regions modulations reported across bilingual infants, children, and adults (Arredondo, Hu, Satterfield, & Kovelman, 2016; Arredondo, Aslin, & Werker, 2021; D’Souza & D’Souza, 2016), that also depend on task complexity (Davis, Kragel, Madden, & Cabeza, 2011; Davis & Cabeza, 2015). Surprisingly, bilingualism was associated with reduced anterior CC, despite positive effects on reading, consistent with previous work in this ABCD sample showing that favorable perinatal conditions were linked to prolonged CC development in presence of better cognitive outcomes (Wu, et al., 2025). This potentially protracted CC development may stem from an increased need to rely simultaneously on opposing inhibitory and excitatory callosal forces (see (Bloom & Hynd, 2005; Knaap & Ham, 2011)) imposed by complex bilingual environments. Indeed, recurrent reliance on effortful, top-down cooperative strategies between left and right frontal regions in bilinguals could effectively go “against” functional lateralisation principles that rely on interhemispheric inhibition, and may slightly protract anterior CC development. However, over time, these strategies may become more automated and more readily deployed at a lower cost, enhancing processing efficiency and supporting the resolution of difficulties, thereby manifesting as a benefit rather than a detriment. This could explain why a positive mediation path from bilingualism to reading was observed, despite appearing negative before controlling for brain size.

Functionally, these adaptive frontal strategies might reflect the reliance on high-order oral language skills (such as morphosyntactic predictions, see Klimovich-Gray et al., 2026) which may protect against family risks of dyslexia and their associated phonological difficulties (Snowling M. J., 2008). It remains unclear whether and how bilingualism-induced anterior CC effects could also influence posterior callosal connections (see (Ronderos, Zuk, Hernandez, & Vaughn, 2024)), directly linked to phonological and reading development (Dougherty, et al., 2007; Swanson, et al., 2015) and protective factors in dyslexia (Yu, et al., 2020).

An important remaining question is when hemispheric rebalance becomes beneficial for reading development and whether positive effects emerge differently for bilinguals with language pairs varying on phonological distance. Here, we showed that bilingual hemispheric rebalancing supports reading in a sample mainly composed of languages with highly distinct phonemic repertoires (inferred Spanish-English bilingualism: over 80% for children and over 73% for parents), but similar effects were observed several times using a behavioral index of hemispheric rebalance (dichotic listening) in Basque-Spanish bilinguals with highly similar phonological repertoires (Ershaid, 2026; Lallier, Peréz-Navarro, & Ordin, 2024). This suggests benefits across language pairs and cultural contexts, although future research should examine these differences more directly.

### 3. Beyond the CC as the only mediator of the link between bilingualism and reading

It is important to stress that anterior CC volume is unlikely to be the primary pathway through which bilingualism influences reading. Indeed, the direct effect of bilingualism on reading was robust across all specifications and substantially larger than the indirect pathway through the anterior CC. Accordingly, in the SEM, there was a limited variance in CC volume explained by the predictors and covariates, consistent with the implication of other structures.

Intrahemispheric connectivity might play a particularly important role in this respect, as bilingual experience has been repeatedly linked to changes in intrinsic functional and structural intrahemispheric connectivity (Fedeli, Del Maschio, Sulpizio, Rothman, & Abutalebi, 2021; Sulpizio, Del Maschio, Del Mauro, Fedeli, & Abutalebi, 2020; Luk, Bialystok, Craik, & Grady, 2011; Pliatsikas, Moschopoulou, & Saddy, 2015; Schlegel, Rudelson, & Tse, 2012; Mohades, et al., 2012; Hämäläinen, Sairanen, Leminen, & Lehtonen, 2017; Singh, et al., 2017). Lower reading skills have also been associated with reduced LH asymmetry of the FA of the arcuate fascicus (AF) - a white matter tract part of the superior longitudinal fasciculus (SLF) linking temporo-parietal to frontal regions of the reading and attentional control networks (Meisler & Gabrieli, 2022; Thiebaut de Schotten, et al., 2011; Vandermosten, Poelmans, Sunaert, Ghesquière, & Wouters, 2013; Zhao, Thiebaut de Schotten, Altarelli, Dubois, & Ramus, 2016) - asymmetry that might be influenced by the CC itself (Andrulyte, Demirkan, Branzi, Bonnett, & Keller, 2026). We initially explored the correlation between the FA asymmetry index of the whole SLF, as provided by the ABCD database (including SLF I, II, III, and AF) (Janelle, Iorio-Morin, D’amour, & Fortin, 2022), and reading skills, but found no significant relationship (see Figures S2,S3). As this lack of correlation could not be attributed to the asymmetry of specific subsegments, we did not pursue further analyses. Future research should further explore the role of the AF/SLF, but also investigate how subcortical and cortical grey matter structures implicated in dyslexia (e.g., IFG, cerebellum) may be modulated by bilingualism (Marin-Marin, Costumero, Ávila, & Pliatsikas, 2022; Pliatsikas, et al., 2020) to influence reading trajectories.

### 3.4 Conclusion

This study offers a nuanced mechanistic understanding of how early bilingual exposure interacts with genetic predispositions to shape the brain’s resilience to reading difficulties. Contrary to the persistent myth that multilingualism exacerbates reading difficulties, our findings demonstrate that bilingual environments may provide a protective, independent buffer against genetic risks for dyslexia through altering interhemispheric connectivity. While genetic predispositions primarily influence reading outcomes through global brain size, bilingualism seems to operate through a distinct neuroanatomical pathway: the anterior CC. Ultimately, these results validate bilingualism as a powerful environmental factor that can foster neurocognitive resilience. We speculate that these frontal structural effects contribute to a more efficient allocation of normally effortful attentional resources across hemispheres, thereby supporting phonological processing and mitigating reading difficulties. These findings open potential avenues for research to test whether leveraging this environmental linguistic factor through educational policies could promote resilience to reading difficulties across the entire ability spectrum.

## 4. Author contributions: CRediT

Conceptualization: M.L., A.C.-C., Data curation: C.R., 23andMe, A.C.-C, Methodology: M.L., A.C.-C, Formal analysis: C.R., A.C.-C. Funding acquisition: A.C.-C., M.L. Supervision: M.L., A.C.-C. Visualization: C.R., A.C.C. Writing – original draft: M.L., C.R., A.C.C. Writing – review & editing: M.L., C.R., A.C.C.

## 5. Author information

### 23andMe research team

Adam Auton, Alan Kwong, Anjali J. Shastri, Barry Hicks, Catherine H. Weldon, David A. Hinds, Emily DelloRusso, Emily M. Rios, Joyce Y. Tung, Kahsaia de Brito, Katelyn Kukar Bond, Keng-Han Lin, Matthew H. McIntyre, Matthew J. Kmiecik, Qiaojuan Jane Su, Robert K. Bell, Sayantan Das, Shubham Saini, Stella Aslibekyan, Vinh Tran, Wanwan Xu, Alisa P. Lehman, Noura S. Abul-Husn, R. Ryanne Wu, Rebecca M. K. Berns, Ruth I. Tennen, Stacey B. Detweiler, Aditya Ambati, Anna Guan, Bertram L. Koelsch, Chris German, Éadaoin Harney, Ethan M. Jewett, G. David Poznik, James R. Ashenhurst, Jingran Wen, Peter R. Wilton, Steven J. Micheletti, and William A. Freyman.

## Acknowledgments

A. C-C. received funding from the Spanish Ministry of Science and Innovation and the Agencia Estatal de Investigación through Ayudas Ramón y Cajal (RYC2022-035511-I). M.L. is supported by the Spanish Ministry of Science and Innovation (grant no. PID2022-136989OB-I00 and ERC-2024-COG 101170337 BIBALANCE). The BCBL acknowledges funding from the Basque Government through the BERC 2022-2025 program, and by the Spanish State Research Agency through The BCBL Severo Ochoa excellence accreditation, CEX2020-001010-S. The funders had no role in study design, data collection and analysis, decision to publish or preparation of the manuscript. We thank the research participants and employees of 23andMe Research Institute. for making this work possible.

Data used in the preparation of this article were obtained from the ABCD^Ⓡ^ Study (https://abcdstudy.org/) and are held in the National Institute of Mental Health (NIMH) Data Archive. This is a multisite, longitudinal study designed to recruit more than 10,000 children aged 9–10 and follow them over 10 years into early adulthood. The ABCD^Ⓡ^ Study is supported by the NIH and additional federal partners under award numbers U01DA041022, U01DA041028, U01DA041048, U01DA041089, U01DA041106, U01DA041117, U01DA041120, U01DA041134, U01DA041148, U01DA041156, U01DA041174, U24DA041123 and U24DA041147. A full list of supporters is available at https://abcdstudy.org/federal-partners/. A listing of participating sites and a complete listing of the study investigators can be found at https://abcdstudy.org/principal-investigators/.

## 7. Code availability

The custom code associated with this study is publicly available at https://git.bcbl.eu/ENDD/MS-reading-IHC-ABCD/.

## 8. Data availability

ABCD^Ⓡ^ data are publicly available through the National Institute of Mental Health (NIHM) Data Archive (https://data-archive.nimh.nih.gov/abcd). GWAS summary statistics used in this study are available from the NHGRI-EBI GWAS Catalog https://www.ebi.ac.uk/gwas/downloads/summary-statistics). Dyslexia GWAS summary statistics can be requested from 23andMe Research Institute. (https://research.23andme.com/collaborate/#dataset-access) and are available in accordance with the scientific review and data transfer agreement.

